# Domain gain or loss in fungal chitinases drives ecological specialization toward antagonism or immune suppression

**DOI:** 10.1101/2025.06.16.659886

**Authors:** Ruben Eichfeld, Asmamaw B. Endeshaw, Margareta J. Hellmann, Bruno M. Moerschbacher, Alga Zuccaro

## Abstract

The evolutionary origins of fungal effector proteins remain poorly understood, particularly how structural changes reprogram antimicrobial enzymes into host-adapted immune suppressors. Here, we show that domain modularity drives ecological specialization in chitinase effectors of the beneficial root endophyte *Serendipita indica*. The GH18 chitinase *Si*CHIT, which carries a C-terminal carbohydrate-binding module (CBM5), is expressed during fungal competition and antagonizes the fungal pathogen *Bipolaris sorokiniana* in the rhizosphere, thereby protecting plant roots. Deletion of the CBM5 abolishes this antifungal activity, while fusion of CBM5 to the CBM5-lacking paralog *Si*CHIT2 restores pathogen inhibition. In contrast, *Si*CHIT2 is induced during root colonization and suppresses chitin-triggered reactive oxygen species production, promoting immune evasion and host compatibility. These results identify CBM5 as a modular determinant of effector function, switching chitinase activity between microbial antagonism and host immune suppression. Our findings support an evolutionary scenario in which effector function in planta arises through domain loss and transcriptional divergence from an antimicrobial precursor, consistent with transitions along the saprotrophy-to-symbiosis continuum.

**Significance Statement:** Effector proteins play key roles in shaping fungal interactions with both plant hosts and microbial competitors, yet how these functions evolve remains unclear. Here, we show that structural domain modularity enables ecological specialization of two paralogous chitinases arising from gene duplication in the root endophyte *Serendipita indica*. Through domain deletion and fusion, we demonstrate that a carbohydrate-binding module (CBM5) determines whether a chitinase functions in microbial antagonism or immune evasion. Our findings provide mechanistic evidence for effector diversification via domain loss and transcriptional divergence, supporting an evolutionary trajectory from antimicrobial activity to host adaptation. This work advances our understanding of how modular architecture drives effector evolution and niche specialization in symbiotic fungi.

**Highlights:**

- Gain or loss of a CBM5 binding domain drives effector specialization between fungal antagonism and immune suppression
- CBM5 acts as a modular determinant enabling antifungal activity in the GH18 chitinase *Si*CHIT
- The CBM5-lacking paralog *Si*CHIT2 suppresses host immunity and promotes root colonization
- Functional divergence following gene duplication illustrates evolutionary repurposing of an antimicrobial enzyme into an immune-suppressive effector

## Introduction

Mutualistic fungi are ecologically important members of the rhizosphere microbiome, where they support plant health and influence microbial community structure and function (Martin & Tan, 2025) This diverse group includes ectomycorrhizal and arbuscular mycorrhizal fungi, as well as beneficial root endophytes. Unlike mycorrhizal fungi, root endophytes often colonize plant roots without forming specialized symbiotic structures and, depending on the host and environmental context, can enhance plant performance by promoting nutrient uptake, water acquisition, and tolerance to environmental stress (Vandenkoornhuyse *et al*., 2002; Dastogeer *et al*., 2018; Mesny *et al*., 2021; Mahdi *et al*., 2022; Andreo-Jimenez *et al*., 2023; Zhang *et al*., 2024). These fungi exemplify the evolutionary plasticity of fungal lifestyles, flexibly occupying ecological niches along the saprotrophy-to-symbiosis continuum. To colonize plant roots and shape their niche, beneficial root endophytes must overcome two key challenges: evasion of host immunity and competition with microbial antagonists and plant pathogens, a process recently shown to be supported by synergistic interactions with bacterial members of the core microbiota (Mahdi *et al*., 2022). Both immune evasion and microbial competition are mediated by secreted effector proteins. Effector-mediated immune evasion often involves the degradation or masking of microbe-associated molecular patterns (MAMPs), such as β-1,3/1,6-glucans and chitin, to prevent recognition by host pattern recognition receptors (Lahrmann & Zuccaro, 2012; Fesel & Zuccaro, 2016; Wawra *et al*., 2016). In parallel, effectors can alter the microbiota composition by suppressing microbial competitors, a strategy well documented in plant pathogens (Snelders *et al*., 2020; Snelders *et al*., 2021) and increasingly recognized in mutualistic fungi.

The beneficial root endophyte *Serendipita indica* (*Si*), a member of the widespread fungal order Sebacinales (Basidiomycota), colonizes a broad range of host plants and is known for its ability to promote growth and suppress plant immunity while protecting the host from pathogens. Colonization is confined to the rhizoplane, epidermis, and cortex, and depends on an intact host immune system (Lahrmann *et al*., 2015). In addition to its host-associated functions, *Si* exhibits strong antifungal activity in the rhizosphere, particularly against the pathogen *Bipolaris sorokiniana (Bs)*, the causal agent of spot blotch and root rot in cereals (Sarkar *et al*., 2019; Mahdi *et al*., 2022; Eichfeld *et al*., 2024). This antagonism is mediated by a GH18-CBM5 endochitinase (*Si*CHIT), whose catalytic activity restricts *Bs* growth and recapitulates the protective effect of *Si* in planta (Eichfeld *et al*., 2024).

GH18 chitinases, found across all fungal phyla, cleave β-1,4-glycosidic bonds in chitin, a major structural component of fungal cell walls. These enzymes fulfill diverse roles in nutrient scavenging, hyphal remodeling, and fungal antagonism (Seidl, 2008; Chen *et al*., 2020). Their catalytic activity depends on a conserved DxxDxDxE motif, which defines the substrate-assisted catalytic mechanism characteristic of this family (van Aalten *et al*., 2001; Bußwinkel *et al*., 2018). While some chitinase-derived effectors have lost enzymatic activity and evolved to sequester MAMPs (Fiorin *et al*., 2018), others retain catalytic function and are functionally enhanced by accessory domains such as carbohydrate-binding modules (CBMs) (Limón *et al*., 2001; Limón *et al*., 2004; Monge *et al*., 2018). These non-catalytic domains increase substrate affinity and specificity, particularly for crystalline or polymeric forms of chitin (Itoh *et al*., 2006; Matroodi *et al*., 2013). *Si*CHIT features a C-terminal CBM5 domain, a modular architecture found predominantly in bacteria and Basidiomycota but absent in Ascomycota, suggesting a lineage-specific evolutionary adaptation (Chen *et al*., 2020; Eichfeld *et al*., 2024). Here, we investigate the functional significance of CBM5 in *Si*CHIT and examine how domain architecture modulates effector activity, enabling a switch between microbial antagonism and immune evasion.

We show that the CBM5 domain is essential for the antifungal and plant-protective function of *Si*CHIT. This chitinase is expressed during fungal competition and antagonizes the pathogen *Bs* in the rhizosphere, thereby protecting plant roots. In contrast, its closely related paralog *Si*CHIT2 lacks CBM5 and is induced during root colonization. *Si*CHIT2 promotes evasion of chitin-triggered immunity, preventing the generation of host-derived reactive oxygen species. Through domain-swapping experiments, we demonstrate that CBM5 functions as a modular determinant that switches the ecological role of a chitinase from microbial antagonism to host immune suppression. Together, our findings support a model in which effector specialization emerges through gene duplication, domain loss, and transcriptional divergence, reprogramming an antimicrobial enzyme into a host-adapted immune effector.

## Results

### CBM5 enhances binding to crystalline chitin and fungal cell wall without altering product specificity

To investigate the role of the C-terminal CBM5 domain in *Si*CHIT (PIIN_03543), we generated three recombinant variants, all lacking their native signal peptides: full-length *Si*CHIT, a catalytically inactive mutant (*Si*CHIT^E196Q^), and a truncated version lacking the CBM5 domain (*Si*CHIT^-CBM5^), which retained the proline-rich linker. The truncated construct was produced by excluding the final 141 bp from the coding sequence. All recombinant proteins were purified via affinity chromatography and verified by SDS-PAGE (Fig. S1).

We first assessed chitinase activity on insoluble crab shell chitin using thin-layer chromatography (TLC) and by quantifying reducing sugars after 20 hours of incubation (Fig. 1A, B). These assays, along with endpoint measurements on chitin azure (CA) (Fig. 1C), revealed no major differences between full-length *Si*CHIT and *Si*CHIT^-CBM5^. However, time-course experiments on CA showed that the presence of CBM5 enhanced early hydrolysis (Fig. 1D), suggesting that the domain increases the reaction rate during initial substrate engagement.

**Figure 1:**
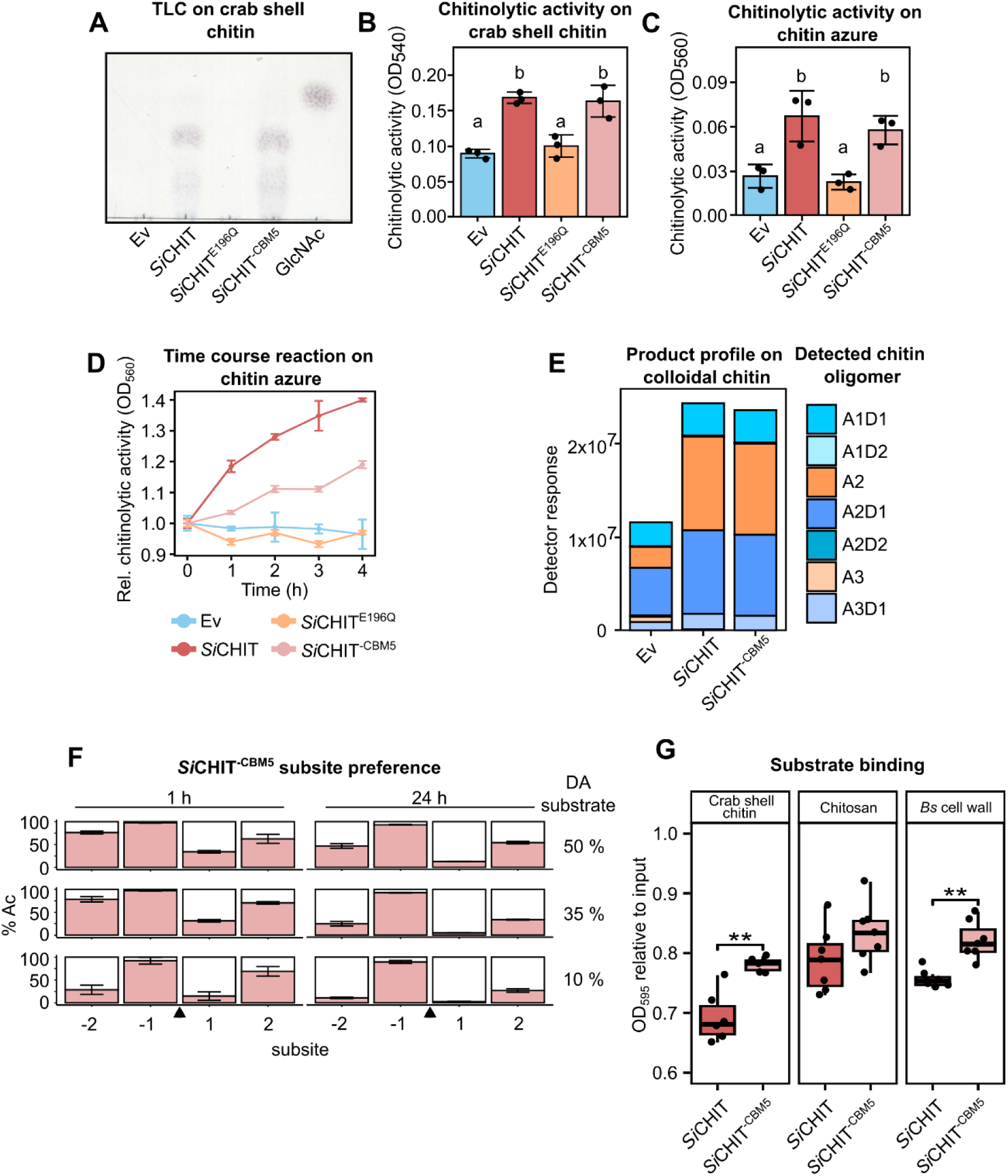
CBM5 enhances binding to fungal cell wall and crystalline chitin and promotes hydrolysis without affecting product specificity. **A)** Thin layer chromatography (TLC) of crab shell chitin incubated with the Ev control or 10 µM of *Si*CHIT, *Si*CHIT^E196Q^, or *Si*CHIT^-CBM5^ at 28 °C for 20 h. **B)** Chitinase activity of recombinant chitinases on crab shell chitin at 10 µM and after 20 h at 28 °C. Samples were spun down and supernatants mixed 1:1 with DNSA reagent. Samples were incubated at 100 °C for 10 min and placed on ice. Subsequently, the relative amount of reducing ends was measured at 540 nm (mean +/- SD, n = 3). **C)** Chitinase activity of recombinant chitinases at 10 µM after 20 h at 28 °C on chitin azure (CA). Samples were spun down and supernatants were quantified at 560 nm (mean +/- SD, n = 3). **D)** Chitinase activity of recombinant chitinases at 10 µM measured over 4 h at 28 °C and normalized to the value of t_0_ (mean +/- SEM, n = 3). **E)** Product profile of *Si*CHIT and *Si*CHIT^-CBM^ on colloidal chitin. Samples were incubated with 5 µM of chitinases for 24 h at 28 °C and hydrolysis products were identified and quantified via MS. A = acetylated, D = de-acetylated unit. **F)** Subsite preference of *Si*CHIT^-CBM5^ measured by MS. Chitosan of three degrees of acetylation (DA) were hydrolyzed for 1 h or 24 h. Based on the sequenced products, the frequency of acetylated units at the −2 to +2 subsites of *Si*CHIT^-CBM5^ was determined. The black arrow indicates the gylcosidic bond between the −1 and +1 subsite that is cleaved by the enzyme. **G)** Substrate-binding ability of *Si*CHIT and *Si*CHIT^-CBM5^ on crab shell chitin, > 75 % de-acetylated chitosan or lyophilized *Bs* mycelium. Binding was assessed by measuring the protein amount in the supernatant of reactions compared to input amount (n = 6-8). Limits of the boxplots represent the 25th–75th percentile, the horizontal line represents the median and the whiskers the minimum/maximum values without outliers. Statistical analysis: **B** and **D)** Different letters indicate significant differences according to one-way ANOVA followed by Tukey’s HSD test (adjusted p-value < 0.05). G) Student’s t-test (P-value: * < 0.05; ** < 0.01).

To evaluate whether CBM5 influences product outcome, we analyzed the hydrolysates of colloidal chitin by mass spectrometry. Both *Si*CHIT and *Si*CHIT^-CBM5^ produced similar oligomer profiles dominated by chitobiose (A2), indicating that CBM5 does not alter product specificity (Fig. 1E). Notably, substantial amounts of partially deacetylated products, such as A1D1 and A2D2, were also detected. This likely reflects the fact that colloidal chitin is not fully acetylated; deacetylated regions may form stretches with higher solubility and increased accessibility, leading to their preferential cleavage by the enzymes. To assess whether CBM5 affects subsite preferences, we analyzed the hydrolysis products of chitosans with defined degrees of acetylation (DA). *Si*CHIT^-CBM5^ displayed the same requirement for an acetylated unit at the −1 subsite and no detectable shifts at adjacent positions compared to *Si*CHIT, confirming that CBM5 does not influence catalytic specificity (Fig. 1F; Eichfeld *et al*., 2024).

To determine the role of CBM5 in substrate interaction, we performed pull-down assays using crab shell chitin, highly deacetylated chitosan, and lyophilized, protein-free *Bs* mycelium. *Si*CHIT^-CBM5^ showed reduced binding to crab shell chitin compared to full-length *Si*CHIT, while no difference was observed with chitosan, suggesting that CBM5 enhances binding specifically to acetylated, crystalline substrates (Fig. 1G). Binding to *Bs* mycelium was also reduced in the absence of CBM5, indicating that the domain contributes to cell wall targeting.

Together, these results show that while CBM5 is dispensable for full catalytic activity and does not alter product specificity, it enhances early hydrolysis rates and substrate binding, particularly to crystalline or fungal cell wall-like chitin. This suggests that CBM5 improves functional efficiency under biologically relevant conditions, likely contributing to the role of *Si*CHIT in microbial competition.

### CBM5 is required for antifungal activity and plant protection

We previously demonstrated that *Si*CHIT restricts the growth of *Bs* and reduces its colonization of plant roots through its chitinolytic activity (Eichfeld *et al*., 2024). However, the specific contribution of the CBM5 domain to this antifungal function remained unresolved. To test whether CBM5 is required for *Bs* inhibition, we incubated *Bs* spores with the full-length *Si*CHIT, the catalytically inactive mutant *Si*CHIT^E196Q^, the truncated variant lacking CBM5 *Si*CHIT^-CBM5^, or the empty vector control Ev. As expected, *Bs* spore germination was significantly reduced in the presence of active *Si*CHIT, but not with *Si*CHIT^E196Q^. *Si*CHIT^-CBM5^ showed a similar lack of inhibitory effect as *Si*CHIT^E196Q^ and had only a minimal effect compared to the Ev control (Fig. 2A).

**Figure 2:**
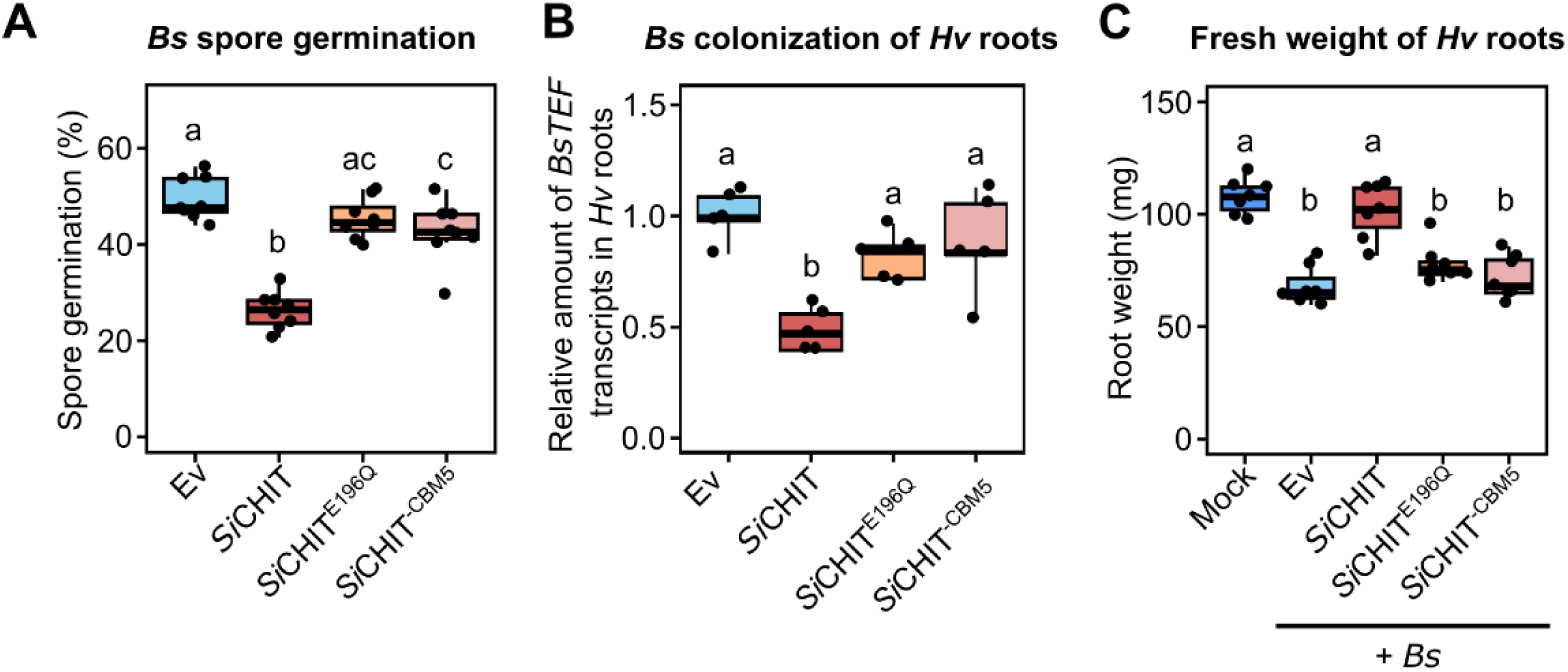
The CBM5 domain is essential for the antifungal and plant-protective activity of SiCHIT. **A)** Relative *Bs* spore germination six h post incubation in sterile chamber slides with the Ev control or 10 µM of *Si*CHIT, *Si*CHIT^E196Q^ or *Si*CHIT^-CBM5^ (n = 6). **B)** Colonization of *Hv* roots by *Bs* at three dpi inferred from relative expression of the fungal housekeeping gene *BsTEF* compared to the *Hv* housekeeping gene *HvUBI* inferred by qPCR using the 2^-ΔCT^ method. Colonization values were normalized to the Ev control. *Bs* spores were either pre-treated with the Ev control or 10 µM of recombinant chitinases for 20 h at 28 °C (mean +/- SD, n = 4). **C)** *Hv* root fresh weight at three dpi with *Bs* spores. *Bs* spores were either pre-treated with the Ev control or 10 µM of recombinant chitinases for 20 h at 28 °C (n = 5-7). Limits of the boxplots represent the 25th–75th percentile, the horizontal line represents the median and the whiskers the minimum/maximum values without outliers. Statistical analysis: Different letters indicate significant differences according to one-way ANOVA followed by Tukey’s HSD test (adjusted p-value < 0.05).

We next examined whether the loss of antifungal activity translates into reduced protection of host roots. *Bs* spores treated with the same chitinase variants were used to inoculate the roots of the *Bs* host *Hordeum vulgare* (*Hv*). As previously shown, *Si*CHIT-treated spores led to a significant reduction in *Bs* colonization, as measured by fungal transcript levels in roots. In contrast, *Si*CHIT^E196Q^ and *Si*CHIT^-CBM5^ had no effect (Fig. 2B). Correspondingly, only *Si*CHIT mitigated the reduction in root biomass caused by *Bs* infection (Fig. 2C). To test whether this effect is conserved in other hosts, we repeated the experiment using *Arabidopsis thaliana* (*At*). *Si*CHIT reduced *Bs* biomass and preserved root elongation, while *Si*CHIT^E196Q^ did not. *Si*CHIT^-CBM5^ had a weak protective effect, albeit significantly lower than the full-length enzyme. (Fig. S2.).

Together, these results demonstrate that both chitinase activity and the CBM5 domain are required for *Si*CHIT’s antifungal and plant-protective functions. The loss of protection observed in the CBM5-lacking variant suggests that this domain contributes to cell wall binding and is critical for effective niche defense.

### A CBM5-lacking paralog, *Si*CHIT2, is transcriptionally specialized for host colonization

GH18-CBM5 chitinases are absent in Ascomycota but occur frequently in the Agaricomycetes (Basidiomycota), where gene copy number varies across species with different ecological lifestyles (Eichfeld *et al*., 2024). To better define the structural requirements for antifungal activity in *Si*, we compared the four GH18 chitinases encoded in the genome with respect to phylogenetic relationships, domain architecture, and transcriptional regulation. Of the four, only *Si*CHIT harbors a CBM5 domain, whereas the other three chitinases contain no additional domains besides the GH18 catalytic domain (Fig. 3A). *Si*CHIT and its closest paralog *Si*CHIT2 (PIIN_03542) share the highest sequence identity and form a distinct phylogenetic pair among the *Si* GH18 chitinases (Fig. 3A; Supplementary Fig. S3). The catalytic DxxDxDxE motif located at the −1 subsite is conserved across all four chitinases, suggesting that each is catalytically active (Fig. 3B). Despite this shared motif, the chitinases exhibit distinct structural features. *Si*CHIT2 retains an 86-amino-acid C-terminal linker but lacks a CBM5 domain. *Si*CHIT3 (PIIN_11727) lacks a predicted signal peptide, while *Si*CHIT4 (PIIN_07603) features extended surface-exposed loops within the GH18 domain (Supplementary Table S1). Taken together, *Si*CHIT and *Si*CHIT2 are more similar to each other in sequence, structure, and genomic context than to the other chitinases in *Si*. Notably, they are located adjacent in the genome (Fig. 3C). Their tandem organization, high sequence identity, and absence of CBM5 in *Si*CHIT2 support a gene duplication event followed by structural divergence through domain loss.

**Figure 3:**
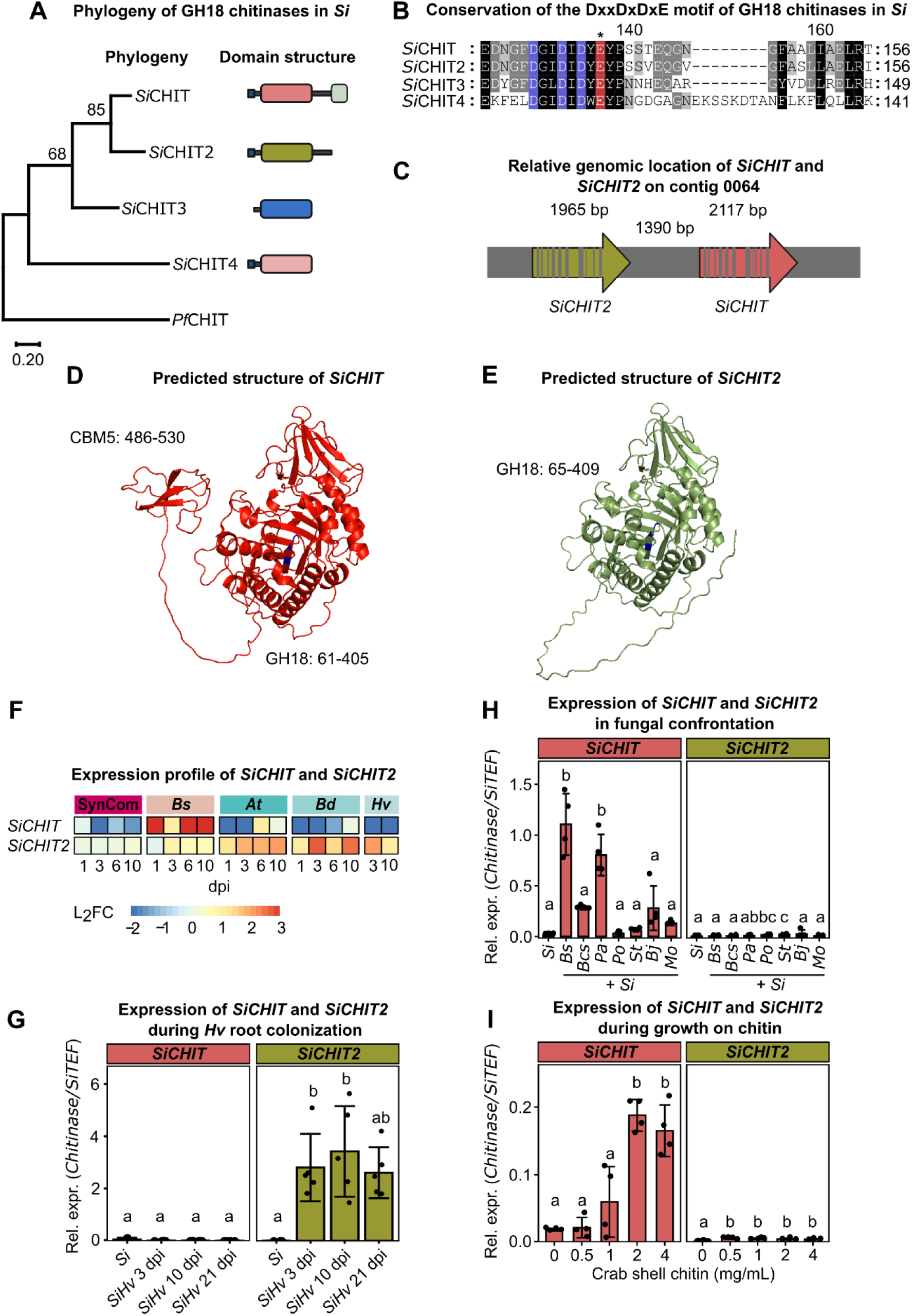
*SiCHIT2* encodes a GH18 chitinase without CBM5 and is expressed during host plant colonization. **A)** Maximum likelihood tree of all four GH18 chitinases encoded by *Si*. Signal peptides were predicted by SignalP 5.0 and removed prior to amino acid sequence alignment. Amino acid sequences were aligned using the MUSCLE algorithm. A maximum likelihood tree was constructed with 500 bootstraps. A GH18 chitinase from *Plasmodium falciparum* was used as outgroup. Symbols on the right indicate the domain architecture of the full length proteins: small blue box: signal peptide; grey line: inter-domain regions; larger boxes (red, green, blue, beige): GH18 domain; small green box in *Si*CHIT: CBM5. Amino acid sequence alignment of the catalytic DxxDxDxE motif. Blue: conserved aspartate (D); red: conserved glutamate (E). The asterisk indicates the glutamate that was mutated to glutamin (Q) in the catalytically inactive *Si*CHIT^E196Q^. **C)** Scheme of the relative location of *SiCHIT* and *SiCHIT2* in the genome of *Si* on contig 0064. The grey bars illustrate introns. *SiCHIT* has 11, and *SiCHIT2* 10 introns. **D, E)** Structure prediction of *Si*CHIT and *Si*CHIT2 by AlphaFold. The sequences of predicted signal peptides were identified with SignalP 5.0 and removed prior to structure prediction. **F)** Expression profile of *SiCHIT* and *SiCHIT2* during confrontation with a bacterial community (SynCom), the pathogen *B. sorokiniana* (*Bs*), or during colonization of the host plants *A. thaliana* (*At*), *B. distachyon* (*Bd*) or *H. vulgare* (*Hv*). The data was taken from Eichfeld *et al*. 2024. **G)** Expression of *SiCHIT* or *SiCHIT2* during *Hv* root colonization at 3, 10 and 21 dpi. Four day old *Hv* roots were inoculated with *Si* and grown on 1/10 PNM medium. Expression was inferred by relative expression of the fungal housekeeping gene *SiTEF* compared to *SiCHIT* or *SiCHIT2* by qPCR using the 2^-ΔCT^ method (mean +/- SD, n = 3-5). **H)** Expression of *SiCHIT* or *SiCHIT2* at six days of growth on 1/10 PNM medium with different fungal species (*Bs* = *Bipolaris sorokiniana*; *Bcs* = *Brunneochlamydosporium sp.*; *Pa* = *Papulaspora equi*; *Po* = *Podospora sp.*; *St* = *Stachybotrys sp.*; *Bj*: *Bjerkandera sp.*; *Mo* = *Mortierella sp.*. Fungal liquid cultures were grown in CM and washed with water, mixed in a 1:1 ratio and 1 g mycelium was streaked out on plates. Expression was inferred by relative expression of the fungal housekeeping gene *SiTEF* compared to *SiCHIT* or *SiCHIT2* by qPCR using the 2^-ΔCT^ method (mean +/- SD, n = 4). **I)** Expression of *SiCHIT* or *SiCHIT2* at six days of growth on crab shell chitin-containing PNM medium at different concentrations. *Si* liquid culture grown in CM was washed with water and 1 g mycelium was streaked out on 1/10 PNM plates. Expression was inferred by relative expression of the fungal housekeeping gene *SiTEF* compared to *SiCHIT* or *SiCHIT2* by qPCR using the 2^- ΔCT^ method (n = 4). Statistical analysis: Statistical analysis: Different letters indicate significant differences according to one-way ANOVA followed by Tukey’s HSD test (adjusted p-value < 0.05).

To assess structural similarity, we generated AlphaFold models (Jumper *et al*. 2021; Mirdita *et al*. 2022) of both enzymes after removing the predicted signal peptides. The models confirmed the overall structural conservation of the GH18 catalytic domain and the presence of a C-terminal linker in *Si*CHIT2, but no CBM5 domain (Fig. 3D & E).

To examine transcriptional regulation, we analyzed RNA-seq data from Eichfeld *et al*. 2024 and found that *Si*CHIT and *Si*CHIT2 were the only chitinases significantly upregulated in biotic contexts. *Si*CHIT was specifically induced during fungal confrontation, while *Si*CHIT2 was induced during colonization of barley, *Brachypodium* and *Arabidopsis*. In contrast, *Si*CHIT3 and *Si*CHIT4 were not transcriptionally activated under any tested biotic conditions, including interactions with pathogenic fungi, beneficial bacteria, or different plant hosts (Fig. 3F & G). To extend the RNA-seq findings, we measured the expression of *Si*CHIT and *Si*CHIT2 by qPCR following confrontation with *Bs* and six additional filamentous fungi representing the major fungal lineages Ascomycota, Basidiomycota, and Mucoromycota. *Si*CHIT was induced in several of these interactions, whereas *Si*CHIT2 expression remained low under all tested microbial conditions (Fig. 3H). Additionally, both *Si*CHIT and *Si*CHIT2 were induced when *S. indica* was grown on crab shell chitin as the sole carbon source (Fig. 3I).

Together, these findings suggest that *Si*CHIT2 evolved a distinct function from *Si*CHIT following gene duplication. Their divergent domain architecture and expression profiles indicate functional specialization, with *Si*CHIT adapted for fungal antagonism and *Si*CHIT2 for host colonization.

### Fusion of CBM5 reprograms *Si*CHIT2 for fungal antagonism

To test whether the CBM5 domain of *Si*CHIT is sufficient to confer antifungal activity, we constructed a chimeric protein consisting of the GH18 catalytic domain of *Si*CHIT2 fused to the proline-rich linker and CBM5 domain of *Si*CHIT (*Si*CHIT2^+CBM5^). As a control, we generated a second construct in which only the linker was fused to the GH18 domain of *Si*CHIT2 (*Si*CHIT2^+link^) (Fig. 4A). As observed for native *Si*CHIT and *Si*CHIT^-CBM5^, the presence of CBM5 did not affect endpoint chitinase activity on crab shell chitin or chitin azure (Fig. 4B, C). However, only *Si*CHIT2^+CBM5^ suppressed *Bs* spore germination, comparable to the effect of full-length *Si*CHIT. In contrast, *Si*CHIT2^+link^ showed no inhibitory activity (Fig. 4D). Consistent with these results, both *Si*CHIT and *Si*CHIT2^+CBM5^ reduced root growth inhibition in barley caused by *Bs*, and limited pathogen colonization. In contrast, *Si*CHIT^-CBM5^ and *Si*CHIT2^+link^ had no measurable protective effect (Fig. 4E & F).

**Figure 4:**
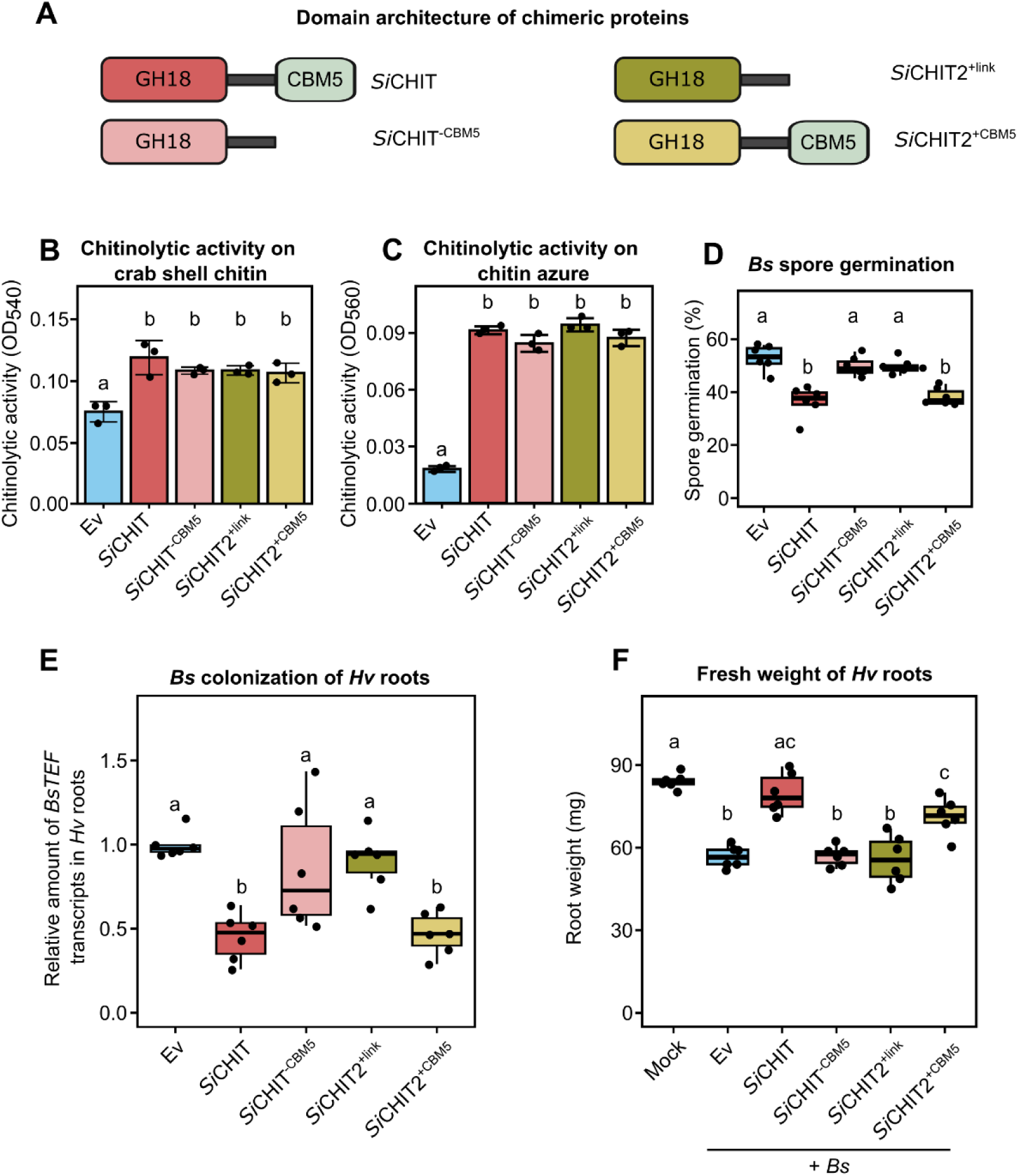
Fusion of the CBM5 to *Si*CHIT2 activates its antifungal and plant-protective capacity. **A)** Domain architecture of recombinant proteins. *Si*CHIT2^+link^ contains the GH18 domain of *Si*CHIT2 and the linker region of *Si*CHIT. *Si*CHIT2^+CBM5^ contains the GH18 domain of *Si*CHIT2 and the linker region with the CBM5 of *Si*CHIT. **B)** Chitinase activity of recombinant chitinases on crab shell chitin at 10 µM and after 20 h at 28 °C. Samples were spun down and supernatants mixed 1:1 with DNSA reagent. Samples were incubated at 100 °C for 10 min and placed on ice. Subsequently, the relative amount of reducing ends was measured at 540 nm (mean +/- SD, n = 3). **C)** Chitinase activity of recombinant chitinases at 10 µM after 20 h at 28 °C on chitin azure (CA). Samples were spun down and supernatants were quantified at 560 nm (mean +/- SD, n = 3). **D)** Relative *Bs* spore germination six h post incubation in sterile chamber slides with Ev control or 10 µM of *Si*CHIT, *Si*CHIT^-CBM5^, *Si*CHIT2^+link^ or *Si*CHIT^+CBM5^. (n = 6). **E)** Colonization of *Hv* roots by *Bs* at three dpi inferred from relative expression of the fungal housekeeping gene *BsTEF* compared to the *Hv* housekeeping gene *HvUBI* by qPCR using the 2^-ΔCT^ method. Colonization values were normalized to the Ev control. *Bs* spores were either pre-treated with the Ev control or 10 µM of recombinant chitinases for 20 h at 28 °C (n = 6). **F)** *Hv* root fresh weight at three dpi with *Bs* spores. *Bs* spores were either pre-treated with the Ev control or 10 µM of recombinant chitinases for 20 h at 28 °C. Limits of the boxplots represent the 25th–75th percentile, the horizontal line represents the median and the whiskers the minimum/maximum values without outliers (n = 6). Statistical analysis: Different letters indicate significant differences according to one-way ANOVA followed by Tukey’s HSD test (adjusted p-value < 0.05).

These results demonstrate that fusing CBM5 to *Si*CHIT2 is sufficient to confer antifungal activity, indicating that CBM5 acts as a modular domain enabling functional adaptation toward microbial antagonism.

### *Si*CHIT2 suppresses chitin-triggered immunity and facilitates root colonization by *S. indica*

Given the distinct plant-induced expression profile of *Si*CHIT2, we hypothesized that this chitinase contributes to immune evasion during root colonization. Previous studies have shown that *Podosphaera xanthii* uses chitinase-like effectors to degrade immunogenic chitin oligomers and suppress host defense responses (Martínez-Cruz *et al*., 2021). To test whether *Si* chitinases function similarly, we incubated immunogenic chitohexaose (A6) with recombinant *Si*CHIT, *Si*CHIT2, their respective variants, or the Ev control at a 1:1 molar ratio. The resulting hydrolysates were applied to barley roots to measure reactive oxygen species (ROS) production. ROS accumulation was completely abolished when chitohexaose was pre-treated with either *Si*CHIT2^+link^ or *Si*CHIT^-CBM5^. In contrast, treatment with *Si*CHIT2^+CBM5^ or full-length *Si*CHIT resulted in only partial or no suppression of ROS accumulation (Fig. 5A, B). An intermediate reduction in ROS accumulation was observed when chitohexaose was pre-treated with *Si*CHIT^E196Q^, likely due to the inactive catalytic cleft sequestering a fraction of the chitohexaose. None of the recombinant chitinases induced ROS when applied in the absence of chitohexaose (Fig. S4).

**Figure 5:**
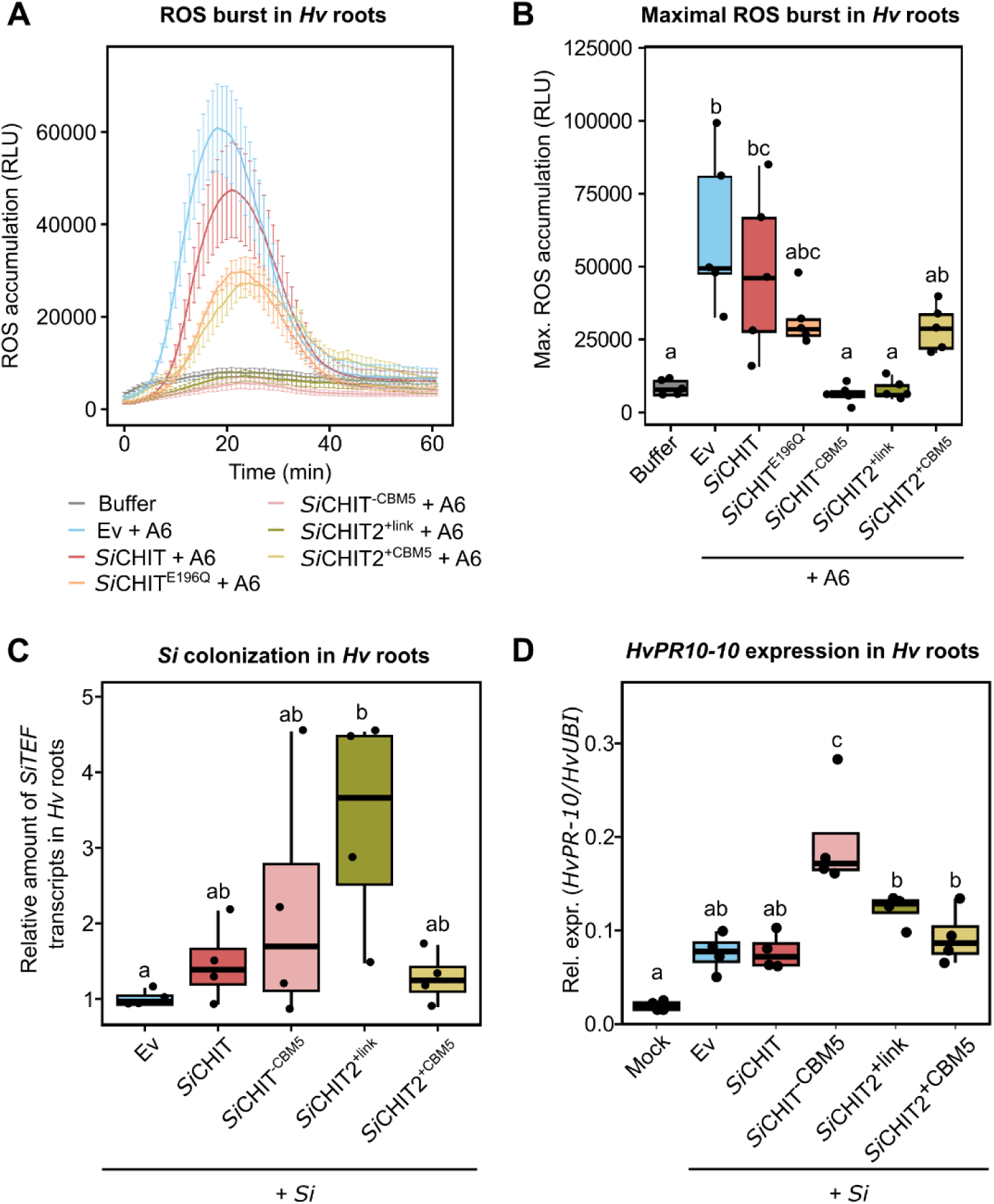
*Si*CHIT2 suppresses chitin-triggered immunity and facilitates root colonization by *Si* in *Hv*. **A)** ROS burst of 4 d *Hv* roots after treatment with 125 nM chitohexaose (A6). Chitohexaose was incubated with the indicated recombinant chitinases in an equimolar concentration (10 µM) 20 h prior to treatment of *Hv* roots. Values represent means +/- SD from five to six individual wells containing four root pieces of ∼ 0.5 cm. **B)** Maximum values of ROS accumulation of *Hv* roots (data from **A**). **C)** Colonization of *Hv* roots by *Si* at three dpi inferred from relative expression of the fungal housekeeping gene *SiTEF* compared to the *Hv* housekeeping gene *HvUBI* by qPCR using the 2^-ΔCT^ method. Colonization values were normalized to the Ev control. *Si* spores were germinated for 16 h at 28 °C and either incubated with the Ev control or 20 µM of recombinant chitinases immediately before *Hv* root inoculation (n = 4). **D)** Transcript accumulation of the *Hv* defense marker gene *Hv*PR10 at three dpi inferred by relative expression of *HvPR10* relative to the *Hv* housekeeping gene *HvUBI* by qPCR using the 2^-ΔCT^ method (n = 4). Limits of the boxplots represent the 25th–75th percentile, the horizontal line represents the median and the whiskers the minimum/maximum values without outliers. Statistical analysis: Different letters indicate significant differences according to one-way ANOVA followed by Tukey’s HSD (adjusted p-value < 0.05).

**Figure 6:**
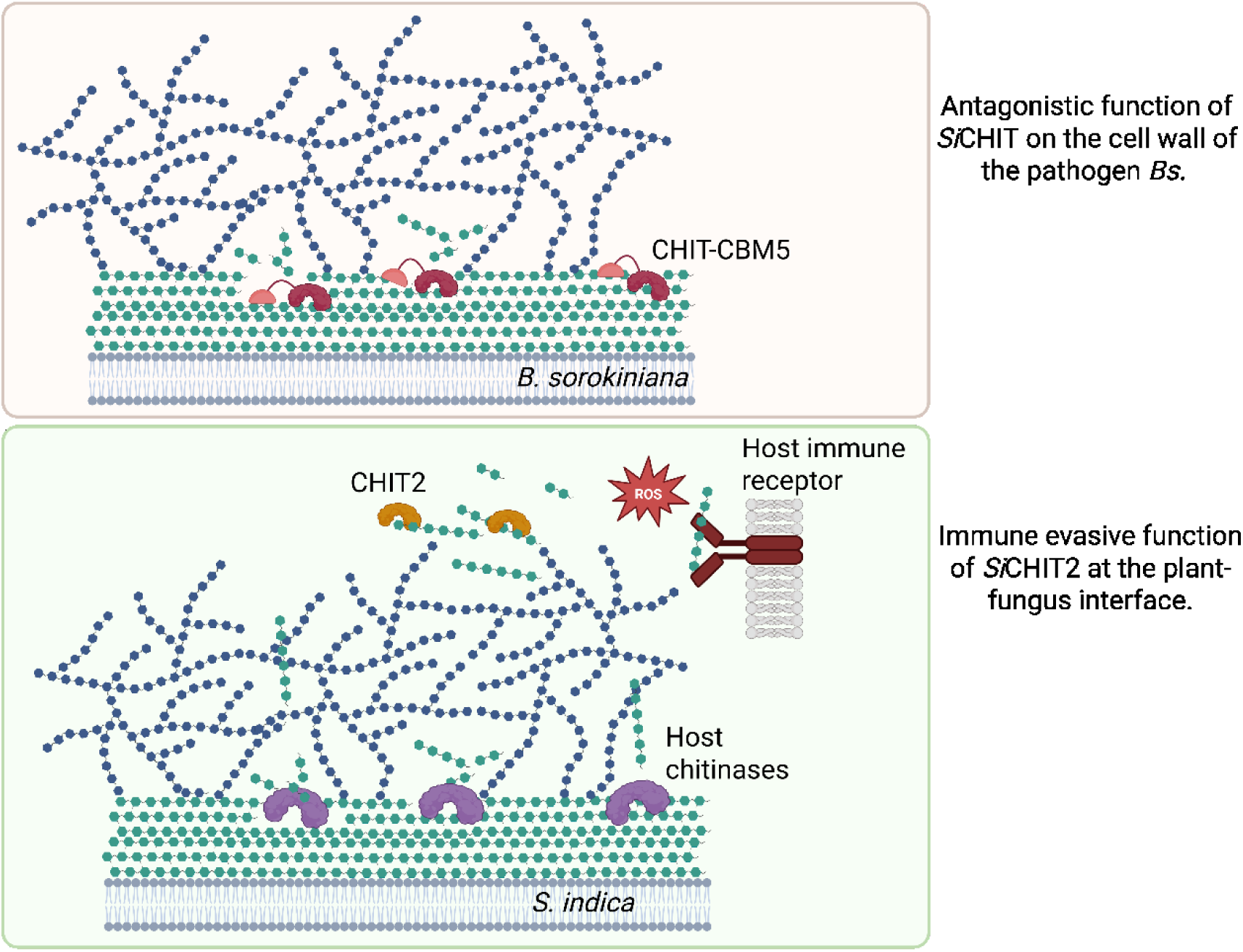
Model displaying the divergent functions of *Si*CHIT and *Si*CHIT2. During confrontation with the pathogen *Bs*, *Si* secretes a GH18-CBM5 chitinase that efficiently inhibits the growth of *Bs*, enabling *Si* to secure its ecological niche. To effectively function in this context, *Si*CHIT relies on the presence of the CBM5. The paralogous *Si*CHIT2 has a catalytically active GH18 domain but no CBM5. It serves *Si* during host colonization by preventing the elicitation of chitin-triggered immunity at the fungus-plant interface, putatively by degrading immunogenic chitin oligomers that are released from the cell wall of *Si* by host chitinases. Figure was created using Biorender.

To assess whether this immune suppression contributes to host colonization, we co-inoculated *Si* with 20 µM of each recombinant chitinase and quantified fungal biomass in barley roots at three days post-inoculation. *Si*CHIT2^+link^ significantly enhanced root colonization, whereas the other chitinases did not (Fig. 5C). *Si*CHIT^-CBM5^ showed a slight but non-significant trend toward increased colonization.

These results suggest that *Si*CHIT2’s ability to suppress ROS depends on the absence of CBM5, which may otherwise restrict access to soluble substrates and favor binding to insoluble chitin. This immune-suppressive function, together with its host-specific transcriptional induction, underscores a distinct role for *Si*CHIT2 compared to its CBM5-containing paralog.

## Discussion

Effector proteins are central to the ecological success of root-colonizing fungi, mediating interactions not only with host plants but also with competing microbes. While traditionally associated with immune suppression (Stergiopoulos & de Wit, 2009), fungal effectors are now also recognized for their roles in inter-microbial competition. They facilitate the acquisition and maintenance of ecological niches in the rhizosphere and on root surfaces, thereby supporting fungal persistence and competitive advantage (Snelders *et al*., 2020; Gómez-Pérez *et al*., 2023; Eichfeld *et al*., 2024). These dual roles are not restricted to pathogens; beneficial fungi such as *Si* employ similar strategies to persist in complex microbial communities, maintain host compatibility, and defend their niche. This functional versatility reflects broader evolutionary trends across the fungal kingdom, where transitions from saprotrophy to symbiosis are accompanied by the repurposing of ancestral traits to meet ecological demands.

Here, we show that *Si* encodes two closely related GH18 chitinases, *Si*CHIT and *Si*CHIT2, that have diverged structurally and transcriptionally to perform distinct functions. *Si*CHIT, which carries a C-terminal CBM5 domain, is expressed during fungal confrontation and contributes to pathogen growth inhibition and plant protection. In contrast, *Si*CHIT2, is induced during root colonization, and suppresses chitin-triggered immune responses in the host (Fig. 5). This divergence possibly reflects an evolutionary trajectory driven by gene duplication and domain loss, enabling *Si* to balance microbial antagonism and host compatibility through modular effector specialization.

The importance of domain modularity in effector function is supported by previous studies in *Trichoderma* spp., where the addition of substrate-binding domains such as CBM1 or CBM18 increases their binding affinity for insoluble substrates and enhances antifungal activity by improving proximity and substrate targeting (Limón *et al*., 2001; Limón *et al*., 2004). However, such effects can be masked *in vivo* by genome-wide redundancy in cell wall-degrading enzymes and antimicrobial effectors. Using purified proteins, we demonstrate that the CBM5 domain is critical for the antifungal activity of *Si*CHIT. Its deletion abolishes fungal antagonism and its reintroduction into *Si*CHIT2 restores this function. This is consistent with studies in bacterial chitinases, where CBM5 loss reduces antifungal activity, likely due to impaired substrate binding (Tsujibo *et al*., 2003; Itoh *et al*., 2006). In our assays, CBM5 enhanced early hydrolysis rates and binding to crystalline chitin and fungal cell wall, but did not affect overall catalytic activity or product profile. This suggests a proximity-based enhancement mechanism, in which CBM5 anchors the enzyme at the fungal cell wall. Unlike antimicrobial or agglutinating lectins such as WSC3, a β-1,3-glucan-binding lectin from *S. indica* that agglutinates fungal cells independently of enzymatic activity (Wawra *et al*., 2019), CBM5 alone does not provide antifungal activity which requires a catalytically active GH18 core. Some fungal pathogens employ catalytically inactive chitinase effectors (e.g., *Mp*Chi in *Moniliophthora perniciosa*; Fiorin *et al*., 2018 or active chitinases (e.g., *Mo*Chia1 in *Magnaporthe oryzae*; Yang & Yu, 2019; or EWCAs; Martínez-Cruz *et al*., 2021) to suppress host immunity by sequestering or degrading free chitin oligomers. Similarly, we show that both *Si*CHIT2 and *Si*CHIT^-CBM5^ can suppress chitohexaose-induced ROS production. These findings suggest that CBM5 may hinder activity on soluble substrates not by altering catalytic specificity, but by limiting access to short oligomers through spatial constraints. CBM5 domains are structurally optimized for engaging extended, crystalline chitin surfaces via aromatic stacking and hydrogen bonding (Kezuka *et al*., 2006; Vaaje-Kolstad *et al*., 2019). Accordingly, CBM5-containing chitinases like *Si*CHIT likely localize to insoluble polysaccharides in fungal cell walls, anchoring the enzyme to non-immunogenic material. Although such substrate targeting could in principle restrict access to soluble MAMPs, this is unlikely in the ROS assay, where only chitohexaose is present. A more plausible explanation is that CBM5 increases the rigidity or steric bulk of the protein, thereby reducing flexibility and limiting productive interaction with short ligands. In contrast, CBM5-lacking variants such as *Si*CHIT2 remain free to access and degrade free immunogenic chitin oligomers. This suggests that suppression of host immunity may require a CBM5-free configuration optimized for apoplastic MAMP degradation during root colonization. Such structural neofunctionalization provides a plausible trajectory for the evolution of fungal effector functions. Given the saprotrophic ancestry and facultative symbiotic lifestyle of *S. indica*, *Si*CHIT likely evolved under selective pressure from fungal competitors, whereas *Si*CHIT2 emerged as an immune-suppressive effector during the transition to a root-associated niche. This represents a clear case of functional divergence driven by domain loss and regulatory rewiring. By comparison, the *Verticillium dahliae* effector *Vd*AMP3 illustrates a different evolutionary strategy: co-opted from an ancestral antimicrobial protein, it retains both its antimicrobial activity and core structural features while functioning in microbiota manipulation in planta. Rather than undergoing major neofunctionalization, *Vd*AMP3 shows how effectors can be repurposed for new ecological roles with minimal structural change (Snelders *et al*., 2021). This highlights the diversity of evolutionary pathways by which fungal effectors adapt to changing environments.

Together, our findings show that domain gain and loss can reprogram effector proteins for distinct roles. CBM5 serves as a modular determinant, switching GH18 chitinase activity between microbial antagonism and immune suppression. This highlights how structural modularity enables fungi to adapt effectors to the dual pressures of rhizosphere competition and host interaction in symbiotic fungi.

## Material and Methods

### Plant and fungal materials

*Hordeum vulgare* (*Hv*, L. cv Golden Promise) and *Arabidopsis thaliana* (*At*, Col-0) were used as plant hosts. *Hv* seeds were surface-sterilized in 6 % sodium hypochlorite for 1 h under constant shaking and subsequently washed five times for 30 min in sterile water. Seeds were germinated for four days on wet filter paper at 21 °C in the dark. Seeds of *At* were surface-sterilized in 70 % ethanol for 10 min and 100 % ethanol for 7 min. *At* seeds were placed on ½ MS medium (Murashige-Skoog medium including vitamins, pH 5.6) containing 1 % sucrose and stratified for three days in the dark and 4 °C prior to germination on a day-night cycle of 8/16 h at 22/18 °C, 60 % humidity and 125 µmol/m^2^ light intensity for seven days. *Serendipita indica* (*Si*, DSM11827) and *Bipolaris sorokiniana* (*Bs*, ND90Pr) were used as fungal species. *Si* was grown on CM medium (Complete Medium, 1.5 % agar, Hilbert *et al*. 2012) at 28 °C for 28 days in the dark. *Si* spores were harvested by adding sterile water and scraping the spores with a scalpel. The suspension was filtered through a sterile miracloth filter and spores were centrifuged at 2,000 x g. The supernatant was discarded and spores were washed three times in water. *Bs* was grown on modified CM medium (Sarkar *et al*., 2019) at 28 °C for 14 days in the dark. *Bs* spores were harvested like *Si* spores but a Drigalski spatula was used to scrape the spores instead of a scalpel.

### Preparation of fungal inoculants and plant colonization

For the preparation of *Bs* inoculants, spores were harvested from 21-day old plates and incubated with the respective recombinant proteins (10 µM in 50 mM phosphate buffer) for 16 h at 28 °C in the dark. For the preparation of *Si* inoculants, spores were harvested from 28-day old plates and were allowed to germinate overnight in sterile water at 28 °C. Prior to plant root inoculation, the respective recombinant proteins were added to 20 µM in 50 mM phosphate buffer. Spore mixtures were diluted to 5000 spores/mL for *Bs* and 500,000 spores/mL for *Si* to yield the final working inoculants. For inoculation of *At*, ten sterile seven-day old seedlings were placed on solid 1/10 PNM (pH 5.6) plates (0.005 % KNO_3_, 0.005 % KH_2_PO_4_, 0.0025 % K_2_HPO_4_, 0.049 % MgSO_4_, 0.00472 % Ca(NO_3_)_2_, 0.0025 % NaCl, 0.5 % (v/v) Fe-EDTA stock solution, 1.2 % Gelrite, pH 5.6. After autoclaving, 10 mL 1 M MES pH 5.6 was added. The EDTA stock solution contains 2.78 % FeSO_4_ x 7 H_2_O, 4.13 % Na_2_EDTA x 2H_2_O). Concentrations are indicated in w/v if not stated otherwise. The roots were treated with 1 mL of the respective inoculant and grown on a day-night cycle of 8/16 h at 22/18 °C, 60 % humidity and 125 µmol/m^2^ light intensity for three days. The root elongation was quantified by measuring the difference root length at the start of the inoculation and the end of the experiment at three days post inoculation (dpi). Roots were washed with water and snap frozen in liquid nitrogen for further RNA extraction. For inoculation of *Hv*, four sterile seedlings were placed in glass jars containing solid 1/10 PNM medium (pH 5.6) and inoculated with three mL of the respective inoculant. The inoculated plants were grown on a day-night cycle of 16/8 h at 22/18 °C and 60 % humidity and a light intensity of 108 µmol/m^2^ for three days. The root weight was quantified at three dpi. Roots were washed in water and snap frozen for further RNA extraction.

### Preparation of fungal confrontations

For the preparation of fungal inoculants, liquid CM medium was inoculated with 500,000 spores/mL. For all other fungi, mycelium was scraped from plates and transferred to liquid CM. All fungi were grown for six days at 28 °C and 120 rpm. The mycelium was blended and re-suspended in fresh CM (Hilbert *et al*. 2012) and regenerated for another 24 h. Regenerated mycelium was filtered through miracloth filters and washed thoroughly with sterile milliQ water. Fungal mycelia were mixed in a 1:1 ratio and 1 g of mycelium was streaked on PNM plates containing 15 % agar. After six days, the mycelia were harvested, dried with tissue papers and snap frozen in liquid nitrogen.

### Fungal spore germination

To quantify the *Bs* spore germination, isolated spores were resolved in liquid CM to a concentration of 100,000 spores/mL and 100 µL were added to the wells of sterile Lab-Tek 2 chamber slides with cover (VWR). 100 µL of the respective treatment (Ev control or purified protein) were added to a final concentration of 10 µM. The germination rate of spores was quantified 6 h after treatment.

### RNA extraction from plant roots and qPCR

Plant materials were ground in liquid nitrogen and 1 mL of TRIzol reagent (Thermo Fischer scientific) was added. Samples were vortexed for 20 sec until all material was resolved. 200 µL chloroform were added and samples vortexed again. Samples were centrifuged at 17,000 x g and 4 °C for 30 min and 500 µL of the upper aqueous phase were transferred to 500 µL isopropanol. The nucleic acids were precipitated at −20 °C for 16 h and pelleted by centrifugation at 17,000 x g and 4 °C for 30 min. Pellets were washed three times with 70 % ethanol diluted in DEPC water. Remaining DNA was removed by adding 1 µL DNase1 (1 U/µL) in 3 µL DNase1 buffer (10 x) (Thermo Fischer scientific) and 26 µL RNase-free water. Samples were incubated at 37 °C for 30 min. DNase digest was stopped by heating to 70 °C for 5 min and subsequent precipitation with 2 volumes isopropanol and ½ volume of 7.5 M ammonium acetate for 16 h at −20 °C. Samples were washed twice with 70 % ethanol in DEPC water and resolved in 30 µL RNase-free water. Concentrations were determined using a NanoDrop 2000c spectrophotometer (Thermo Fischer Scientific).

1 µg of RNA was used for reverse transcription into cDNA (Thermo Fischer Scientific). First, 1 µL of random hexamer and of oligo-dt oligomers were added and incubated at 65 °C for 5 min and subsequently placed on ice. 4 µL 5 x reaction buffer, 1 µL Riboblock RNase inhibitor (20 U/µL), 2 µL 10 mM dNTPs and 2 µL MMLV reverse transcriptase were added and samples were incubated at 42 °C for 1 h. cDNA synthesis was stopped by heating to 70 °C for 5 min. qPCR was conducted using the primers listed in Supplementary Table S2.

### Cloning of chitinase expression vectors

The coding sequence (CDS) of the chitinase genes were amplified by PCR with Q5 High-Fidelity Polymerase (New England Biolabs) using the primers listed in Supplementary Table S2. PCR products were cloned using the Gibson assembly cloning technique into the PQE80L expression vector (Qiagen, Hilden, Germany) after vector linearization with Kpn1 (New England Biolabs).

### Production and purification of recombinant protein in *E. coli*

Chemically competent *E. coli* BL21 cells were transformed with the vectors for protein expression under control of a lac operon. 5 mL starter cultures were prepared in liquid LB medium containing 100 µg/mL carbenicillin and grown for 16 h at 37 °C and 180 rpm. Starter cultures were transferred to 300 mL of LB medium with carbenicillin and grown to an OD_600_ of 0.5 at 28 °C. Gene expression was induced by adding IPTG to a final concentration of 1 mM and cells were grown at 16 °C for 20 h. Cells were pelleted by centrifugation at 5000 g for 15 min at 4 °C. Bacterial pellets were resolved in 5 mL lysis buffer (50 mM NaH_2_PO_4_, 300 mM NaCl and 10 mM imidazole, pH 8) and sonicated three times for 1 min with pulses on ice and centrifuged at 16,200 g for 30 min. 5 mL of the supernatant were added to 1 mL of Nickel-NTA Agarose (Qiagen, Hilden, Germany) and incubated under continuous shaking at 4 °C for 1 h. Samples were spun down at 500 g for 10 secs, the flow through was removed, 4 mL wash buffer (50 mM NaH_2_PO_4_, 300 mM NaCl and 20 mM imidazole, pH 8) were added and samples were incubated under continuous shaking at 4 °C for 10 min. Washing steps were repeated five times. For elution, 1 mL of elution buffer (50 mM NaH_2_PO_4_, 300 mM NaCl and 250 mM imidazole, pH 8) was added and samples were incubated under continuous shaking at 4 °C for 10 min. Eluted fractions were dialyzed against 50 mM phosphate buffer (pH 6) over night and protein concentrations were measured using Bradford reagent. To test the purity of the protein samples, aliquots were boiled for 5 min in SDS-sample buffer, loaded on a 10 % SDS gel and visualized by Coomassie staining.

### Chitinase activity assays

To quantify the enzymatic activity of the purified chitinases, 2 mg/mL chitin azure (Sigma Aldrich) were incubated with 5 µM recombinant chitinase in a total volume of 200 µL. Reactions were incubated at 28 °C for 20 h. Reactions were stopped by heating to 95 °C for 5 min and samples were centrifuged 5 min at 17,000 x g and the absorbance of supernatants was quantified at 560 nm. For quantifying reducing ends, 10 g crab shell chitin were incubated with 5 µM enzyme in 200 µL 50 mM phosphate buffer for 20 h at 28 °C. Reactions were stopped by heating to 95 °C for 5 min. Samples were centrifuced for 5 min at 17,000 x g and supernatant were mixed in a 1:1 ratio with DNSA (3,5-dinitrosalicylic acid) reagent. DNSA reagent is prepared by mixing 1 % (w/v) 3,5-dinitroslicylic acid, 1.6 % (w/v) sodium hydroxide, 30 % (w/v) potassium sodium tartrate tetrahydrate. The samples were heated for 10 min at 95 °C and subsequently transferred to ice. The absorbance of the samples was measured at 540 nm.

For TLC, 10 mg crab shell chitin (Sigma Aldrich) were weighed in 2 mL reaction tubes and recombinant chitinases were added to a final concentration of 5 µM in 200 µL. Reactions were incubated at 28 °C for 20 h and stopped by heating to 95 °C for 5 min. Samples were centrifuged at 17,000 x g for 5 min. 10 µL of the supernatant were separated on 60 F_254_ TLC (Sigma Aldrich) silica gel plates with running buffer (n-butanol/methanol/28 % ammonium hydroxide/water, 5/4/2/1). Plates were dried and then bathed in developer solution (1.6 g diphenylamine/1.6 mL aniline/12 mL 85 % phosphoric acid/80 mL acetone). TLC plates were developed in an incubation oven at 100 °C for 20 to 30 min.

### Chitinase subsite specificity

The subsite specificity of *Si*CHIT^-CBM5^ was determined as previously reported (Cord-Landwehr *et al*., 2017; Eichfeld *et al*., 2024).

### Substrate binding assay

15 mg of crab shell chitin (Sigma Aldrich), Chitosan (Sigma Aldrich) or freeze-dried mycelium of *Bs* were weighed in a 2 mL reaction tube and incubated with 400 µL chitin-binding buffer (2 mM KH_2_PO_4_, 8 mM Na_2_HPO_4_, 2 mM KCL) and 100 µL of 20 µM recombinant protein shaking at 4 °C for 1 h. Samples were centrifuged at 17,000 x g for 5 min and 60 µL of the supernatant were mixed with 60 µL Bradford reagent. Protein amounts were quantified at 595 nm and compared to the input amount.

### Quantification of reactive oxygen species (ROS)

For elicitor preparation, 10 µM of chitohexaose were incubated with 10 µM of recombinant chitinase or the buffer control for 16 h at 28 °C. Reactions were stopped by heating to 95 °C for 5 min and chitohexaose was diluted to 250 nM.

Four root pieces (0.5 cm) of four-day old barley seedlings were transferred to each well of a 96-well microtiter plate (white, flat bottom) containing 200 μL of 2.5 mM 2-(N-morpholino)ethanesulfonic acid (MES) buffer, pH 5.6. The plate was incubated for 16 h at 21°C for recovery. The buffer was replaced with 100 μL 2.5 mM MES buffer containing 20 μM LO-12 and 20 μg/mL Horseradish peroxidase (HRP). After 25-min incubation in the dark, 100 μL of 250 nM chitohexaose elicitor solution was added to each well. Chemiluminescence was measured using a TECAN SPARK 10 M microplate reader over all wells for 2 h with an integration time of 450 ms.

## Data availability

All data supporting the findings of this study are available within the article and supporting information.

## Acknowledgements

We acknowledge support from the Cluster of Excellence on Plant Sciences (CEPLAS) funded by the Deutsche Forschungsgemeinschaft (DFG, German Research Foundation) under Germany’s Excellence Strategy-EXC 2048/1-Project ID: 390686111 and the ZU 263/11-2 (SPP 2125 DECRyPT).

## Author contributions

AZ and RE conceptualized the study. RE and AE conducted most of the research. MJH, BMM and RE conducted mass spectrometric analysis of the product profile and the subsite specificity. AZ and RE wrote the manuscript with input from all authors.

