## Supplementary information for "Domain gain or loss in fungal chitinases drives ecological specialization toward antagonism or immune suppression"

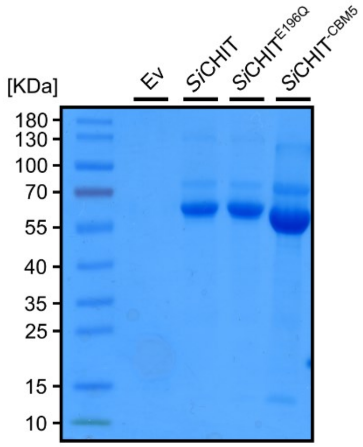


**Figure S1: Purification of chitinases.** The coding sequences with an N-terminal His-Tag were cloned into an expression vector for *E. coli* (PQE80L) and induced with IPTG. The purification was conducted using Nickel-NTA slurry and purified proteins were visualized on a 10 % SDS-Gel and stained with Coomassie brilliant blue.

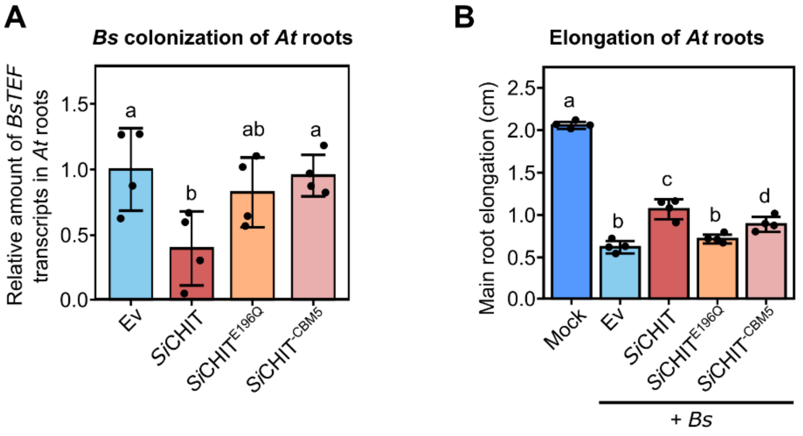


**Figure S2: The CBM5 is essential to protect *A. thaliana* from the pathogen *Bs*. A)** Colonization of *At* roots by *Bs* at three dpi inferred from relative expression of the fungal housekeeping gene *BsTEF* compared to the *At* housekeeping gene *AtUBI* by qPCR using the $2^{-\Delta CT}$ method. Colonization values were normalized to the Ev control. *Bs* spores were either pre-treated with the Ev control or 10 µM of recombinant chitinases for 20 h at 28 °C (mean +/- SD, n = 4). **B)** *At* root elongation at three dpi with *Bs* spores. *Bs* spores were either pre-treated with the Ev control or 10 µM of recombinant chitinases (mean +/- SD, n = 4). Statistical analysis: Different letters indicate significant differences according to one-way ANOVA followed by Tukey' honest significant difference test (adjusted p-value < 0.05).

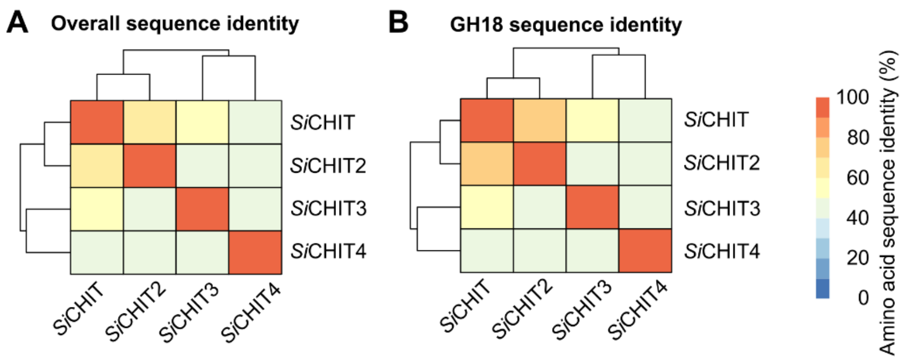


**Figure S3: Sequence identity of the four *Si* GH18 chitinases.** The amino acid sequences without signal peptide were aligned and sequence identity in percent (%) calculated and visualized as heatmap. **A)** Overall sequence identity: the full amino acid sequence without signal peptide was used for the alignment. **B)** GH18 sequence identity: only the amino acid sequence identity of the predicted GH18 domains was used for the alignment.

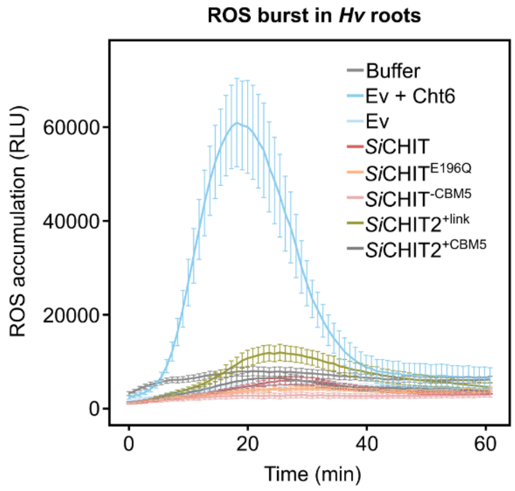


**Figure S4: ROS burst in Hv roots**. ROS burst of 4 d *Hv* roots after treatment with 125 nM chitohexaose. 10 µM chitohexaose (Cht6) were incubated with the Ev control 20 h prior to treatment of *Hv* roots. Roots were treated with the indicated elicitors at 125 nM. Ev + Cht6 and Buffer values from Fig. 5 A were used for proper comparison with the treatments. Values represent means +/- SEM from five to six individual wells containing four root pieces of ~ 0.5 cm.

**Table S1: Domain composition of the four GH18 chitinases from *Si*.**

| Protein ID | No. amino acids | Signal peptide | GH18 domain | CBM5 | Remarks |
|---|---|---|---|---|---|
| PIIN_03543 | 533 | 1-19 | 61-405 | 486-530 | C-terminal CBM5 |
| PIIN_03542 | 495 | 1-16 | 65-409 | None | No CBM5 but C-terminal linker |
| PIIN_11727 | 314 | None | 20-311 | None | No signal peptide |
| PIIN_07603 | 523 | 1-28 | 155-437 | None | Three low complex complexity loops within the GH18<br>a: 154-167<br>b: 286-297<br>c: 406-417 |

**Table S2: Primers used in the study.**

| Primer name | Sequence 5' → 3' | Use |
|---|---|---|
| PIIN_03543_qPCR_for | cctgggtcttgggagaatg | qPCR |
| PIIN_03543_qPCR_rev | ggcgtcgtagcagtatgaa | qPCR |
| PIIN_03542_qPCR_F | acggcatctgggactacaaa | qPCR |
| PIIN_03542_qPCR_R | gcagaaagttcccagtgcat | qPCR |
| HvPR10_FW (Sakar et al 2019) | ggagggcgacaaggtaagtg | qPCR |
| HvPR10_RV (Sakar et al 2019) | cgtccagcctctcgtactct | qPCR |
| Bs_Tef_for (Sakar et al. 2019) | cgccgtaccggaaagtctg | qPCR |
| Bs_Tef_rev (Sakar et al. 2019) | ggcgaaacgaccaagagga | qPCR |
| TEF_Piri_QPCR_F | gcaagttctccgagctcatc | qPCR |
| TEF_Piri_QPCR_R | ccaagtggtgggtactcgtt | qPCR |
| HvUbi60_fwd (Sakar et al. 2019) | accctcgccgactacaacat | qPCR |
| HvUbi60_rev (Sakar et al. 2019) | cagtagtggcggtcgaagtg | qPCR |
| SiCHIT_pQE_FW | atcaccatcaccatcacggatccgcatgcgagctcggtaccacgcccgccatgatgc | cloning |
| SiCHIT_pQE_RV | ctcagctaattaagcttggctgcaggtcgacccggggtacctcagcacgacttgagttgattccac | cloning |
| SiCHITtrun_pQE_RV | ctcagctaattaagcttggctgcaggtcgacccggggtacctcacataccagcacatgttccacc | cloning |
| PIIN_03542_cloning_F | tccgcatgcgagctcggtaccgtcgttggacgtcccaagaa | cloning |
| PIIN_03542_cloning_R | agccccttttgtctgcagaaagttcccag | cloning |
| PIIN_03542_Linker_cloning_F | ttctgcagacaaaaggggctccgagtcg | cloning |
| PIIN_03542_Linker_cloning_R | tgcaggtcgacccggggtacctcacataccagcacatgttccacc | cloning |
| PIIN_03542_Linker_CBM5_F | ttctgcagacaaaaggggctccgagtc | cloning |
| PIIN_03542_Linker_CBM5_R | tgcaggtcgacccggggtacctcagcacgacttgagttgattccacg | cloning |
